# Experimental effects of within-brood genetic variation on parasite resistance in a wild bird host

**DOI:** 10.1101/2022.05.26.493631

**Authors:** Jessica Gutiérrez, Conor C. Taff, Suzannah Tupy, Catherine Goncalves, Sarah A. Knutie

## Abstract

Hosts can differ in parasite susceptibility across individuals, populations, and species. Genetic variation can influence parasite susceptibility by affecting host resistance to parasitism. For example, genetic variation among related individuals, such as within broods of offspring, might be a key factor influencing within-brood resistance to ectoparasitism. The goal of this study was to determine if within-brood variation of eastern bluebirds (*Sialia sialis*) affects susceptibility to ectoparasitic blowflies (*Protocalliphora sialia*). To address the goal, we conducted a partial cross-fostering study for which half of the nestlings were cross-fostered (experimental) or not (control). Nestling physiology (i.e., glucose, hemoglobin, and parasite-specific IgY antibody levels), morphometrics (i.e., mass, tarsus length, bill length, and first primary feather), survival, nestling status (resident versus fostered), and sex were characterized. We also quantified parasite abundance, life stage, and pupal size. We found that experimental nests had fewer parasites and more larvae than pupae compared to control nests, which suggest that within-brood genetic variation affects parasite abundance. However, this effect was driven by the sex ratio with the experimental nests, with female-biased nests having fewer parasites than male-biased nests. Treatment did not affect nestling morphometrics, physiology, or survival at the nest level. Within experimental nests, resident females had significantly higher hemoglobin levels when compared to fostered females. Resident and fostered males and fostered females had significantly higher glucose levels than resident females. Together, these results suggest that resident females were fed upon less than fostered females and may have been less stressed than males and fostered females. Overall, our study demonstrates the importance of considering within-brood variation and nestling sex in understanding host-parasite interactions.

## Introduction

Parasite abundances are seldomly uniform across individuals, populations, and host species (Anderson & May 1982, Holt *et al*. 2003, Lively 2010). Host resistance mechanisms, such as the immune response, can vary among individuals, which might explain variation in parasite abundance (Råberg *et al*. 2009). For example, individuals that mount an antibody (immune) response have fewer parasites than individuals that do not mount this response, resulting in variation in parasite abundance within a population (Råberg *et al*. 2009, Owen *et al*. 2010). Host resistance might be particularly important to early-life host health and survival when young can be “sitting ducks” for parasites. Identifying the factors that influence early-life host resistance, such as genetic diversity (Penn *et al*. 2002, Blanchet *et al*. 2009) and sex (Zuk & McKean 1996, Klein 2002), might be important to explain variation in parasite abundance within a population.

The effect of genetic diversity on host resistance has been considered at the population and individual level. For example, increased host genetic variation, at the population level, may enhance overall resistance to parasites (Altermatt & Ebert 2008, Lively *et al*. 2014, Ekron *et al*. 2019). Individual hosts with increased diversity of major histocompatibility complex (MHC) can increase antigen recognition (Penn *et al*. 2002, Sommer 2005). However, genetic variation within a collective of related individuals, such as a brood, can shape within-brood resistance against parasites (Jennions & Petrie 2000).

Many taxa, including fish, amphibians, mammals, and birds, have mating systems for which broods from a single female are fertilized by multiple males (Hasselquist & Sherman 2001, Rolland *et al*. 2003, Chen *et al*. 2011, Hain and Neff 2007). Most wild bird species, including passerines, are socially monogamous but engage in multi-male mating (most commonly referred to as extra-pair copulation; Hasselquist and Sherman 2001). The benefits of extra pair copulations have been relatively well studied: males that have offspring in multiple nests increase their reproductive success, compared to putting “all of their eggs in one basket”. However, the potential benefits of extra-pair mating for females are not as well understood. One explanation is that by engaging in extra-pair mating, females increase within-brood genetic diversity to increase parasite resistance. Studies that have compared immune metrics between within-and extra-pair young have found varying results. For example, Arct *et al*. (2013) found that differences in the immune responses of within- and extra-pair young depended on environmental conditions. In contrast, Wilk *et al*. (2008) found that the immune response did not differ between within-and extra-pair young. However, studies are often correlational, rely on immunocompetence assays, rather than responses to actual parasites (Owen *et al*. 2010), and focus on individual-level variation rather than brood-level effects.

The goal of this study was to determine whether within-brood genetic diversity affects eastern bluebird (*Sialia sialis*) resistance to parasitic nest flies (*Protocalliphora sialia*). Bluebirds and their parasitic nest flies are an ideal system to understand how genetic diversity can influence host defenses against parasites for several reasons. Bluebirds commonly use artificial nest boxes to raise their offspring, which means that researchers can easily use experimental manipulations, such as cross-fostering experiments, to establish causation. Adult blowflies are non-parasitic but they lay their eggs in the nests of birds, shortly after nestlings have hatched. Once fly eggs hatch, the larvae feed on the blood of nestlings before pupating in the nests and metamorphosing into and eclosing as adult flies (Sabolsky *et al*. 1989). Blowflies feed on the collective brood of nestlings within a nest and overall broods have variable resistance to *P. sialia* (Grab *et al*. 2019, Knutie 2020). Additionally, bluebirds have relatively low occurrence of extra-pair young within broods (Stewart *et al*. 2010) when compared to other passerines.

In our study, we experimentally manipulated within-brood genetic composition using partial-cross fostering of nests (hereon, experimental nests) or not (hereon, control nests). Nestling morphometrics (body mass, tarsus length, feather length), survival, hemoglobin levels (proxy for blood loss), glucose levels (a potential proxy of stress), the *P. sialia*-specific IgY antibody response, and sex were then characterized. We predicted that higher within-brood genetic diversity would increase overall brood resistance to parasitism, and therefore nests with partially cross-fostered nestlings would have lower parasite abundance and disrupted parasite development (less parasite larvae develop into the pupae) when compared to control nests. We also predicted that parasite abundance would be negatively correlated to the mean *P. sialia*-specific antibody response of the nest. In turn, nests with fewer parasites were predicted to have less blood loss (hemoglobin), higher body mass, larger morphometrics, and higher survival. In closely related mountain bluebirds (*Sialia currucoides*), nestling sex affects parasite feeding, with flies consuming larger blood meals from females compared to males (Brien & Dawson 2013). Higher female to male ratios can affect parasite abundance, size, and development. We predicted that eastern bluebird nest sex ratio will affect parasite abundance, life stage of the flies, and pupal size across treatments. Our study will lead to a better understanding of how within brood genetic diversity influences host-parasite interactions in wild populations, as well as the potential benefits of extra-pair copulations for female birds.

## Materials and methods

### Study system

Our study was conducted near the University of Minnesota Itasca Biological Station and Laboratories (47°13’33” N, −95°11’42” W) in Clearwater and Hubbard County. From May through August 2020, we followed 200 wooden nest boxes, which are commonly used by eastern bluebirds. Bluebirds are secondary cavity nesters that readily build and rear offspring in nest boxes. In our focal population, bluebird clutch sizes range from 1-6 eggs, eggs are incubated for 14-16 days, and nestlings will remain in the nest for 15-19 days (Grab *et al*. 2019, Knutie 2020). Blowflies are the primary nest parasite of bluebirds in this study area (Grab *et al*. 2019).

### Partial cross-fostering experiment

Nest boxes were monitored weekly for egg laying by the bluebirds (Fig 1). Once all eggs were laid (determined by a cease in egg-laying after 24 hours), the bluebird mother was left undisturbed for 14 days to incubate the eggs. Afterward, nests were checked for hatching once a day. After the first day of hatching (hatch day = 0), the first two nests (synchronized) were randomly assigned a treatment (control or experimental); the following paired set of nests were then assigned to the opposite group and so on in alternating order. Nests were considered synchronized if hatching of eggs occurred 0-24 hours apart from each other. When all nestlings from a set of treatment nests had hatched (1-2 days after the first nestling hatch), 1-3 nestlings from one nest were cross-fostered or sham-cross-fostered with another nest. Early cross-fostering is necessary to avoid environmental and parental biases that are associated with this method (Winney *et al*. 2015).

**Fig. 1.**
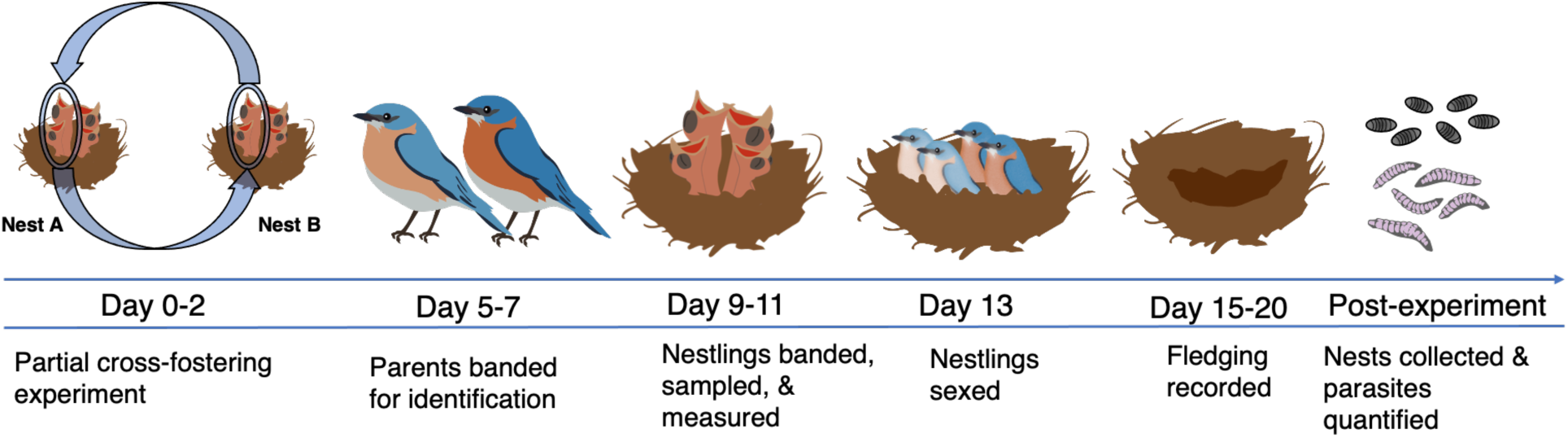
Timeline and experimental design for partial cross fostering study. Graphics by Melissa Ingala.

Prior to partial cross-fostering, all nestlings were weighed to the nearest 0.1 g using the Brifit Digital Mini Scale. The nestlings were then numerically ranked from lightest to heaviest in mass. The heaviest nestlings from both nests were randomly assigned to stay in the original nest or be switched and the following nestlings were assigned in alternate order. This was done to avoid size differences within a nest; see Giordano *et al*. (2014) for cross-fostering details. After nestlings were weighed, the natal down feathers were partially trimmed into unique cuts to identify nestlings at day 10; the location of these trims were as follows: 1) no cut, 2) right or left shoulder, 3) right or left head, and 4) top or bottom back) and 5) a combination of these cuts (i.e., left head and bottom back cut). The cut combinations were assigned a number then the order of cuts was randomly selected at a nest (i.e., cut patterns 1-4 were assigned to one nest, then cut patterns 5-8 were assigned to the synchronized nest to prevent having the same cut patterns within one nest). To prevent artificial parasite introduction to nests, nestlings were checked thoroughly and attached parasites were removed and returned to the original nest. To avoid stress during the transfer, nestlings were kept in a padded box with a Multi-purpose Jumbo 72-hour Uniheat Heat Pack. Time out of the nests did not exceed 35 minutes. Control nestlings were handled using the exact same method but returned to their original nests.

### Quantifying nestling growth and survival

On day 10, nestlings received a Fisheries and Wildlife numbered metal band and a unique color band combination for individual identification after fledging. Nestlings were weighed (to the nearest 0.1 g) using an Ohaus CS200-100 portable compact scale and measured (bill length [0.1 mm], right tarsus length [0.1mm], and left first primary feather [0.1mm]) using analog calipers from Avinet. Since nestling sex can affect feeding preferences of the parasite (O’Brien & Dawson 2013), nestlings were sexed on day 13 using visual differences in feather pigmentation (O’Brien & Dawson 2009). Starting on day 15, nests were checked every other day from a distance to determine the age of fledging. Total nestling survival for a nest was determined by the proportion of nestlings that survived out of the total number of nestlings.

### Nestling and adult tissue collection

During cross-fostering (day 1-2), nestling buccal cells were collected by orally inserting and swirling a sterile thin foam swab on both sides of the buccae. Buccal swabs are a minimally invasive way to collect DNA from nestlings that are too small in mass for blood collection. Buccal swabs provide less genetic material than blood samples; however, collecting these swabs will prevent a loss of sample if the nestlings naturally die before 10 days old (when blood samples are collected). Buccal swabs were placed in a 0.6 mL tube in 300 μL of 70% ethanol, kept in a cooler with ice packs in the field, and stored in a −80°C freezer.

On day 10, after approximately three minutes of banding, a small blood sample (<30 μL) was collected from the nestlings’ brachial vein using a 30-gauge needle and a 70 μL heparinized capillary tube. One to two resident and fostered nestling blood subsamples (>6 μL) were haphazardly chosen to measure glucose (mg/dl) and hemoglobin (g/dl) using the Accu-Chek Performa and the HemoCue® Hb 201+ Portable Analyzer, respectively. The rest of the whole blood was placed in a 0.6 mL snap top tube and then in a cooler with ice packs in the field until we returned to the lab. The tubes were centrifuged at 6,000 x g for 3 minutes to separate the red blood cells from the plasma. The plasma was placed into 0.6 mL tubes and stored at −80°C. The blood cells were removed from the tube and preserved on a Whattman™ FTA Card at room temperature to dry for 24 hr prior to transferring to a −20°C freezer. At the end of the season, all samples were shipped on dry ice to the University of Connecticut; plasma and buccal swabs were stored in a −80°C freezer until used in immunoassay or DNA extractions, respectively. The blood cards were stored at −20°C until used in the DNA extractions.

Adult bluebirds were captured during systematic nest-box monitoring and post-hatching days 5-7. To capture adults, we cut Hygloss Overhead Projector Sheets into a 3.5 × 6.5 cm vertical rectangle and covered the nest box hole from the inside. We stabilized the top side of the cutout with 2 push pins. The cutout was placed 1.5 cm above the top part of the hole, overlapping all other sides by 2 mm. Once an adult bluebird entered the nest box, a black extra-large hooded sweatshirt was placed over the box, we partially opened the door, and carefully reached inside to handle the adult bluebird. If parents were captured, we collected buccal swabs (same as nestling methods) and measurements (mass [0.01 g], wing length [0.5 mm], bill length [0.1 mm], and right tarsus [0.1 mm]). Finally, the bluebirds received a Fisheries and Wildlife numbered metal band and a unique color band combination for individual identification.

### Quantifying parasite abundance and size

Once the nestlings fledged or died (15-21 days), nests were collected and dissected for *P. sialia* larvae (1st, 2nd, or 3rd instars), pupae, and pupal cases to quantify total parasite abundance. If the nest had parasites with pupae, we measured 1-5 pupae and measured the length (0.01 mm) and center width (0.01 mm) with analog calipers. The parasites were separated by life stage and were stored in 50 mL falcon tubes with original nest material and closed with a breathable mesh cap. During nest collection 5-10 3rd instar larva were placed in 2 mL screw cap tubes then stored in −80°C freezer. After 1-3 days of collection, a subset of the pupated larvae was measured for a total of 10 parasites (i.e., 5 pupae were measured at collection and 5 pupae after storing the larvae) and placed back into their tubes. If the nest had no pupae at collection, then a subset of 10 pupated larvae were measured. The length and width measurements were used to calculate pupal volume (V = π*[0.5*width]^2^*length). Finally, 10-14 days after the collection date, 2-3 flies were collected in 90% EtOH in 2 mL screw cap tubes, the remaining were released near the Itasca Biological Station and Laboratories.

### Blood and buccal swab DNA extractions

For a subsample of nests (n = 14), blood and buccal DNA were extracted using the Qiagen DNeasy Blood and Tissue Kit protocol with modifications. Buccal cells and blood DNA were extracted from the swabs and Whatman™ FTA Cards, respectively. Blood cards were hole-punched and the fragments were placed in 1.5 mL microcentrifuge tubes; for buccal swabs we added the swab and 70% EtOH solution into 1.5 mL microcentrifuge tubes. For both sample types, we then added 30 μL of proteinase K, 190 μL 1X PBS, added 200 μL of Buffer AL, and vortexed. Samples were incubated on a heat block at 56°C; samples were vortexed for 15 s every 5 minutes for a total of 35 minutes. After the incubation step, swabs were removed and200 μL of 100% EtOH was added to the sample mixture and then pipetted into the spin column while leaving the filter paper behind. The Qiagen protocol was then followed with no modifications.

### Assigning parentage and quantifying within-brood genetic diversity

Extractions were then followed by polymerase chain reaction (PCR) using the Type-it Microsatellite PCR Kit protocol with modifications of the annealing temperatures. Five fluorescently labeled primers were amplified: Smex9 (Ferree *et al*. 2008), Smex14 (Ferree *et al*. 2008), Pdo5 (Griffith *et al*. 1999), Sialia36 (Faircloth *et al*. 2006), and Sialia37 (Faircloth *et al*. 2006) their annealing temperatures were 65°C, 59°C, 59°C, 59°C and 62°C respectively. Each primer was selected based on previous amplification success using eastern bluebird DNA (Faircloth *et al*. 2006, Ferree *et al*. 2008, and Stewart *et al*. 2009). The total volume per reaction was 12.5 μL: 6.25 μL of 2x Type-it Multiplex PCR Master Mix, 4 μL RNase-free water, 1.25 μL of 10x fluorescently labeled primer mix, and 1 μL of DNA template. The final concentrations for 2x Type-it Multiplex PCR Master Mix,10x primer mix, the DNA template were 1X, 2μM, and 10 ng, respectively. PCR products were analyzed by Life Science SeqStudio Capillary Electrophoresis at the University of Connecticut CORE facilities.

We called the alleles for each microsatellite locus using the ‘Fragman’ v1.0.9 package in R version 4.0.4 and following the workflow described in the package documentation (Covarrubias-Pazaran et al. 2015). We included all adults and nestlings for which we were able to reliably call peaks for at least three of the five loci. After calling peaks, nestlings were compared first to the putative mother. The majority of nestlings matched their mother at each scored locus. In four cases, nestlings mismatched the putative mother at one or more loci. Because we could not be sure if these mismatches resulted from sample collection or genotyping errors, these nestlings were excluded from paternity assignment. For the rest of the nestlings, we compared the remaining alleles to the putative father and considered any nestling that mismatched the putative father at one or more loci to be an extra-pair offspring. Because we did not sample all males in the population, we did not attempt to assign the identity of extra-pair fathers.

### Quantifying the parasite-binding antibody response

Enzyme-linked immunosorbent assays (ELISA) were used to detect *P. sialia*-binding antibody (IgY) levels in nestlings using the Grab *et al*. (2019) and DeSimone *et al*. (2018) protocol. Ninety-six well plates were coated with 100 μL/well of *P. sialia* protein extract (capture antigen) and were 1:100 diluted in carbonate coating buffer (0.05 M, pH 9.6). The plate was then sealed with parafilm and incubated for 1 hr on an orbital table (VWR Microplate VRTX 120V ADV). Plates were then washed and coated with 200 μL/well of bovine serum albumin (BSA) blocking buffer then sealed and incubated for overnight at 4°C. Between each of the following steps, plates were washed with a Tris-buffered saline wash solution using a plate washer (Thermo Fisher Scientific Microplate Washer), loaded as described, and incubated for 1 hr on an orbital table at room temperature. Plasma was 1:100 diluted with sample buffer then wells were loaded with 100 μL/well of individual diluted host plasma in triplicate. Plates were t loaded with 100 μL/well of Goat-αBird-IgG-Heavy and Light Chain HRP (diluted 1:50,000; A140-110P; Bethyl Laboratories). Plates were loaded with 100 μL/well of peroxidase substrate (tetramethyl-benzidine, TMB: Bethyl Laboratories) and incubated for exactly 30 min. Finally, the reaction ceased using 100 μl/well of stop solution (Bethyl Laboratories).

Optical density (OD) was measured with a spectrophotometer (PowerWave HT; 450 nm filter; BioTek). A higher OD value corresponded to higher IgY concentration. On each plate, a positive control of pooled plasma from naturally parasitized nestlings and adults were used in triplicate to correct for interplate variation. We corrected for interplate variation by first dividing the mean OD value for the positive controls for each plate by the highest OD value among all plates then by multiplying the mean for each sample by this correction. In addition, each plate contained two nonspecific binding (NSB1 and NSB2) controls. The NSB1 control contained capture antigen but excluded plasma, while NSB2 contained plasma but excluded capture antigen. Finally, each plate included a blank sample in which only the detection antibody was added, but plasma and capture antigen were excluded. The highest NSB values (NSB1 or NSB2) were subtracted from the mean OD value of each sample to account for binding of the detection antibody to the capture antigen.

### Statistical analyses

All analyses and figures were produced in RStudio (2020, Version 1.2.1335; R Version 3.6.3). A test of normality was conducted on all variables of interest (parasite abundance, parasite larvae to pupa ratio, parasite volume, 1st primary length, tarsus length, mass, hemoglobin, glucose, and antibody levels) using a Shapiro-Wilke test with the Stats package (P > 0.05 was considered normal) (R Core Team). Analyses were conducted using generalized linear models (GLM) and generalized linear mixed models (GLMM) functions with the lme4 package and MASS package (Bates *et al*. 2015; Venables & Ripley, 2002). Nest-level analyses were conducted using a GLM (Poisson) to determine the effects of partial cross-fostering treatment, sex ratio, and its interaction on parasite abundance and nestling age at fledging. A GLM (binomial) was used to determine the effects of partial cross-fostering treatment, sex ratio, and its interaction on nestling survival and parasite life stage (larvae to pupae ratio). Finally, a GLM (family: Gamma, log-link) was used to determine the effects of partial cross-fostering treatment, sex ratio, and its interaction on parasite size (volume).

Nestling morphometrics (mass, bill length, tarsus length, feather length) were not correlated (all R^2^ < 0.70) and therefore, measurements were analyzed using separate GLMM (family: Gamma, log-link) with nests as a random effect. However, tarsus length was found to be normal therefore we used the GLMM family gaussian. A Mann-Whitney T-test was used to analyze the effect of treatment on nestling morphometrics coefficient of variation. A GLMM was used to analyze the effects of cross-fostering treatment on approximately 10-day old nestling physiology (log transformed hemoglobin and antibody levels), with nest as a random effect; glucose levels were found to be normal. A GLMM was also used to analyze the effect of nestling status (resident versus fostered), nestling sex, and interaction on the variables of interest, such as brood size, hatch date, fledging age, or nestling survival. When deciding between which covariates should be used within a model, the model with the lower AICc value was selected. Probability values were calculated using log likelihood ratio tests using the ANOVA type III function in the car package (Fox & Weisberg, 2011). Effect sizes of significant results were analyzed using a Cohen’s d estimate function in the effsize package (Marco Torchiano, 2020).

## Results

### Frequency of extra-pair young

Cross fostering increased the diversity of parental genetic contribution to nestlings. Parentage was assigned to nestlings, for which we had genetic material from both parents, from their original nests (*n* = 14; Table S1). Out of 14 nests, 50% (n = 7) of nests had at least one EPY and nests ranged from 1-4 EPY (20%-100% of the nest). Most nests with EPY (5/7 nests) only had one EPY, whereas the other two nests contained 100% (4/4 nestlings) and 60% (3/5 nestlings) EPY. Overall, 12 out of 44 nestlings (27%) were identified as EPY. Within control nests, all nestlings were genetically related to one mother. Within the control nests for which we assigned parentage to nestlings (n = 8 nests), four nests did not contain EPY, three nests contained one EPY, and one nest contained three EPY. Within experimental nests, all nestlings were genetically related to two mothers. Within the experiment nests (before cross fostering; n = 6), three nests did not contain EPY, two nests contained one EPY, and one nest contained four EPY. Only two pairs of experimental nests (n = 4 nests) were assigned parentage; after cross fostering, all nestlings were assigned to at least two fathers and in one nest, nestlings were assigned to at least three fathers.

### Effect of treatment on parasites

Parasite abundance was significantly affected by treatment, sex ratio, and the interaction between treatment and sex ratio (Fig.2a, Table S2). Parasite abundance was reduced by partially cross-fostering (experimental group) when compared to control (GLM, χ^2^ = 84.92, df = 1, *P* < 0.0001; parasite abundance means: control: 58.8, experimental: 49.9), although, the effect size was small (Cohen’s d estimate: 0.24). The main effect of cross fostering treatment on parasite abundance was likely driven by the effect of the relationship between treatment and sex ratio on parasite abundance (χ^2^ = 22.12, df = 1, *P* < 0.0001); within the experimental group as the proportion of female increased, parasite abundance decreased (Estimate: −0.02) when compared to controls. Sex ratio also significantly affected parasite abundance (χ^2^ = 19.06, df = 1, *P* < 0.0001); as proportion of female increased, parasite abundance decreased (Estimate: −0.48); the effect size of sex ratio was large (Cohen’s d estimate: 1.98). Treatment did not significantly affect the sex ratio of nests (χ^2^ = 0.41, df = 1, *P* = 0.52; proportion of female nestlings: control: 0.51, experimental: 0.45).

Parasite life stage (proportion of larvae to pupae) at fledging was affected by treatment (χ^2^ = 15.15, df = 1, *P* < 0.001, average number of larvae in experimental:16.2 and control: 26.9; Cohen’s d estimate: 0.30) (Fig. 2b). Within the experimental group, the larvae to pupae ratio was lower than compared to the control group (Estimate: −0.87) (Table S2). However, nestling sex ratio alone (χ^2^ = 0.02, df = 1, *P* = 0.88) and the interaction between treatment and sex ratio did not affect parasite life stage (χ^2^ = 0.81, df = 1, *P* = 0.37) (Table S2). Parasite volume was not affected by treatment (GLMM, χ^2^ = 0.94, df = 1, *P* = 0.33), nestling sex ratio (χ^2^ = 0.57, df = 1, *P* = 0.45) or the interaction between the two variables (χ^2^ = 2.60, df = 1, *P* = 0.11) (Fig. 2c).

**Fig. 2.**
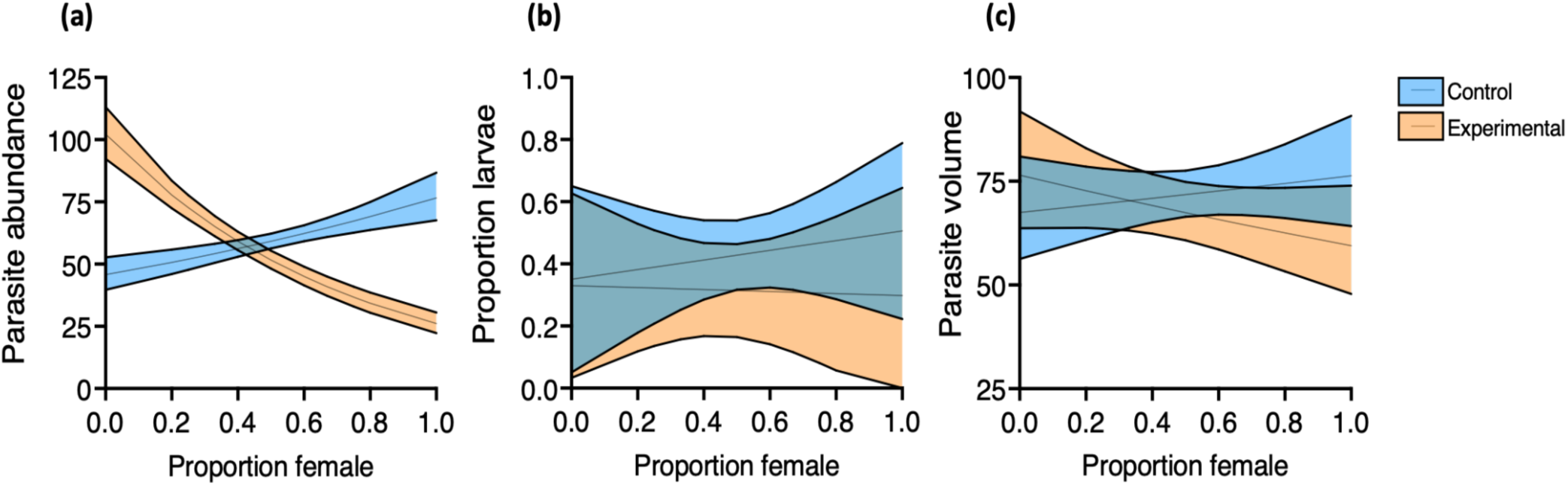
Model results for the effect of the interaction between cross-fostering treatment (control vs. experimental) and sex ratio (proportion of females within a nest) on parasite a) abundance, b) life stage, and c) size. Shaded area represents the standard error of the mean.

### Effect of treatment on nestling survival and health

#### Nestling survival

Nestling survival to fledging was not affected by treatment (GLM, χ^2^ = 3.06, df = 1, *P* = 0.08), nestling sex ratio (χ^2^ = 0.15, df = 1, *P* = 0.70), or the interaction between the two variables (χ^2^ = 1.56, df = 1, *P =* 0.21) (Table S3). Age at fledging was also not affected by treatment (χ^2^ = 0.33, df = 1, *P* = 0.57), sex ratio (χ^2^ = 0.55, df = 1, *P* = 0.46), or interaction between the two variables (χ^2^ = 0.42, df = 1, *P =* 0.52) (Table S3). Within the experimental treatment, nestling survival was not affected by nestling status (resident versus fostered) within the nest (χ^2^ = 0.01, df = 1, *P =* 0.92), nestling sex (χ^2^ = 0.00, df = 1, *P =* 0.99), or the interaction between the two variables (χ^2^ = 0.02, df = 1, *P =* 0.90) (Table S4). Age at fledging within the experimental treatment was also not affected by nestling status (χ^2^ = 0.04, df = 1, *P =* 0.84), nestling sex (χ^2^ = 0.01, df = 1, *P =* 0.93), or the interaction between the two variables (χ^2^ = 0.06, df = 1, *P =* 0.81) (Table S4).

#### Nestling morphometrics

Nestling mass at cross-fostering (0-2 days old) did not differ significantly by treatment (GLMM, χ^2^ = 0.06, df = 1, *P =* 0.80, mean mass in grams for experimental: 4.23 and control: 4.63). At approximately 10 days old, nestling mass, tarsus length, first primary feather length, and bill length were not affected by treatment, sex, or their interaction (Table S5). Similarly, for fostered versus resident nestlings in the experimental treatment, morphometrics were not affected by nestling status, sex, or interaction (nestling status and sex) (Table S6). Treatment also did not affect the coefficient of variation for morphometrics (Mann-Whitney T-test P > 0.05).

### Effect of treatment on nestling physiology

#### Nestling IgY antibodies levels

Nestling IgY antibodies at approximately 10 days old were not affected by sex (χ^2^ = 7.20, df = 1, *P* = 0.71). IgY antibody levels were not affected by treatment (χ^2^ = 1.22, df = 1, *P* = 0.27), or the interaction between the two variables (χ^2^ = 1.02, df = 1, *P =* 0.31) (Table S7-S8). Antibody levels within the experimental group were also not affected by nestling status (χ^2^ = 0.08, df = 1, *P* = 0.78), sex (χ^2^ = 0.47, df = 1, *P* = 0.49), or the interaction between the two variables (χ^2^ = 0.03, df = 1, *P* = 0.86) (Table S8-S9).

#### Hemoglobin

Nestling hemoglobin levels at approximately day 10 were not affected by treatment (χ^2^ = 0.62, df = 1, *P* = 0.43), sex (χ^2^ = 0.28, df = 1, *P* = 0.59), or the interaction between the two variables (χ^2^ = 0.05, df = 1, *P* = 0.83) (Table S7-S8). Within the experimental group, hemoglobin levels were affected by status (χ^2^ = 8.28, df = 1, *P* < 0.004) and the interaction between status and sex (χ^2^ = 11.21, df = 1, *P* < 0.001) (Table S8-S9). However, sex alone did not affect hemoglobin levels (χ^2^ = 2.19, df = 1, *P* = 0.14). Resident female nestlings had significantly higher hemoglobin levels than fostered females, but there were no differences in hemoglobin levels between males and females (Fig 3a).

**Fig. 3.**
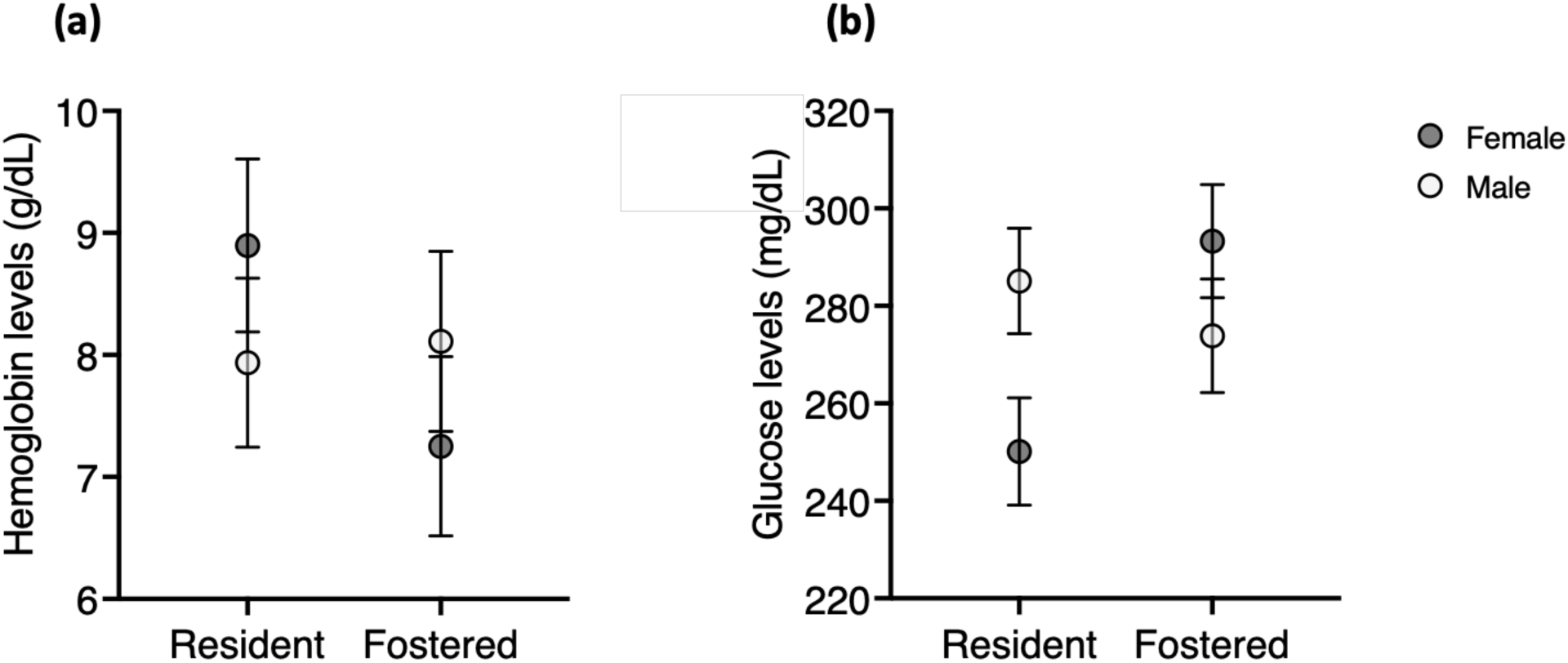
Model results for the effect of the interaction between nestling status (resident versus cross-fostered) and sex on a) hemoglobin levels b) glucose levels. Results represented by the mean +/-standard error.

#### Glucose

At approximately 10 days old nestling glucose levels were not affected by treatment (χ^2^ = 1.39, df = 1, *P* = 0.24), sex (χ^2^ = 0.57 df = 1, *P* = 0.45), or the interaction between the two variables (χ^2^ = 1.56, df = 1, *P =* 0.21) (Table S7-S8). Glucose levels within the experimental treatment were affected by nestling status (χ^2^ = 18.07, df = 1, *P* < 0.0001) and the interaction between status and sex (χ^2^ = 11.21, df = 1, *P* < 0.0001) (Table S8-S9). Glucose levels were not affected by sex alone (χ^2^ = 2.19, df = 1, *P* = 0.14) (Table S8-S9). Resident females had significantly lower glucose levels than fostered females and males (Fig. 3b). However, there were no significant differences between males (resident and cross fostered) or fostered females (Fig. 3b).

## Discussion

We studied the experimental effects of within-brood genetic variation on host resistance to blowflies. Cross-fostered nests had fewer parasites and more larvae (compared to pupae) than the control nests. However, this effect was not driven by the IgY antibody response, suggesting that another immune mechanism is mediating this relationship. Interestingly, the main effect of cross-fostering treatment on parasite abundance was dependent on its interaction with sex ratio, with decreased parasite abundance as the ratio of females within a nest increased. Among the cross-fostered nestlings, fostered female nestlings, and all resident and fostered males had more blood loss and elevated glucose levels compared to resident female nestlings. Our results suggest that the effect of treatment is potentially driven by the effect of cross-fostering on female physiology which may be affected by differing pre- and post-hatching environments. For example, mothers can adjust their offspring’s phenotype to their local environment via altering egg investment and/or nestling provisioning (Coster *et al*. 2012).

We found that experimental nests had fewer blowflies and a higher larvae to pupae ratio compared to control nests. This result suggests that nests with increased within-brood genetic diversity might have lower susceptibility to parasite infestation. Increased within-brood genetic diversity could increase overall resistance, such as via the immune response, to parasitism. For example, Arct *et al*. (2013) increased brood size and thus within-brood genetics and found that extra-pair offspring had increased immune competence. Within a few days after hatching, altricial nestlings can produce an endogenous innate immune response (Pihlaja *et al*. 2006, Palacios *et al*. 2009) and use maternally transferred antibodies (obtained during egg development) to resist parasites (King *et al*. 2010). The immune response can vary among individual nestlings based on their condition, as well as genotype, and in turn affect brood susceptibility to parasites (Boulinier *et al*. 1997, Brinkhof *et al*. 1999, Arct *et al*. 2013). Within our experimental treatment, cross-fostered nestlings had different mothers and fathers, which could have led to changes in genetically-influenced immunological resistance to parasites. Future studies could determine whether changes in nestling resistance was related to paternal or maternal genetics or the combination of the two variables.

The antibody-mediated immune response was not responsible for the observed effect of treatment on parasite abundance in bluebirds. Previous studies have found that bluebird nestlings do not mount a significant IgY antibody response to parasitism when food resources are more restricted (Knutie 2020). Our results suggest that overall, nestlings are likely food restricted since they did not produce a significant IgY antibody response to parasitism. Since the immune response to ectoparasites is complex, other mechanisms of immunity could be responsible for the differences observed in parasitism across treatments (Owen *et al*. 2010). For example, when dermal tissue is damaged by parasite feeding and foreign antigens, such as salivary proteins, are released. The antigens then trigger cytokine release that stimulate phagocytic cell activity. The phagocytic cells migrate to the damaged tissue, engulf the antigens and present fragmented antigens to helper T lymphocytes which leads to T-cell specific antibody production and release. The T and B lymphocytes (memory cells) recognize the antigen after re-exposure which leads to the recruitment of phagocytic and granulocytic cells that causes tissue swelling. Finally, the inflamed tissue can disrupt and prevent parasite feeding. We did not characterize other potential immune metrics, such as cytokine levels, phagocytic cell counts, or oxidative stress and antioxidant activity, but future studies should target these measures to determine the mechanism by which within-brood genetic diversity affects parasitism.

Since control nests have more parasites than experimental nests, blood loss and glucose (a proxy for stress) should be greater in control nests compared to experimental nests. Although mean hemoglobin levels were higher and glucose levels were lower in experimental nests when compared to control nests this difference was not significant. Parasite abundance was statistically higher in control nests, but the mean number of parasites in control nests was 58.8 compared to 49.9 in experimental nests, which is a mean difference of 8.9 parasites; if the difference in parasite abundance had been more extreme, then we might expect to see significantly higher hemoglobin and lower glucose levels within the experimental nests.

The interaction between treatment and sex also affected parasite abundance; within the experimental group, parasite abundance decreased in female-biased nests compared to male-biased nests. These results suggest that female-biased nests are less susceptible to parasitism as compared to male-biased within the experimental treatment. In conflict with our own findings, O’Brien and Dawson (2013) found more blowflies in female biased mountain bluebird nests compared to male-biased nests, which suggests that females are more susceptible to parasitism than males. However, female nestlings can decrease oxidative stress and increase antioxidant activity when exposed to parasites (Coster *et al*. 2012). Increased antioxidant activity can reduce free radicals inside a cell and thus allow damaged tissue to heal faster. Our result suggests that females may exhibit increased resistance to parasites than males, but further investigation is needed to determine the mechanism.

Although male-biased experimental nests had more parasites than female-biased experimental nests, hemoglobin did not differ significantly between males and females. One potential explanation is that blowflies have been found to take smaller blood meals from males than females (O’Brien & Dawson 2013). Another explanation is that there are sex differences in blood recovery, but more research is required to understand these differences in developing nestlings (Murphy 2014). Within the experimental nests, fostered and resident males had more blood loss (although non-significant) and higher glucose levels than resident females. However, fostered females had similar hemoglobin and glucose levels compared to males suggesting that fostered females were more stressed and fed upon than resident females. The differential stress responses between sexes across treatments could be responsible for our observed result, but additional studies are needed to determine causation.

The act of experimentally moving nestlings across nests changes the biological and physical environment of developing nestlings (Mateo & Holmes 2004). Brood manipulation can introduce unintended biases such as differences in nestling growth, survival, and nest sex ratio (Winney *et al*. 2015). However, we found no such effect of treatment on nestling morphometrics, age at fledging, sex ratio, or survival which is similar to Giordano *et al*. (2014) partial cross-fostering study. The results of our study show that although parasite abundance differed between treatment, the effect of these interactions on the factors measured are marginal.

Individual host variation can create within-brood susceptibility to parasitism that leads to variation in parasite abundance (O’Brien & Dawson 2013). The results of this study may be important in understanding host-parasite interactions and how this shapes parasite and host population structure. Our hypothesis that within-brood variation and relatedness affect parasite metrics (parasite abundance and life stage), because overall individual variation can lead to over nest resistance was supported. Furthermore, our hypothesis that within-brood variation would indirectly affect nestling physiology, development, and fledging since parasitism can affect blood metrics, immune response, and eventually nestling development was not supported. How individual and population level genetic variation in animals affects host-parasitism remains an open question (Lively *et al*. 2014); however, our study demonstrates that future research may need to consider within-brood and litter variation to determine its effect on host-parasite interactions and life histories.

## Land Acknowledgement

The authors and collaborators acknowledge the Ojibwe people who have cared for and occupy the land in which our research is conducted. The University of Minnesota Itasca Biological Station and Laboratories is located on the land ceded by the Mississippi and Pillager Bands of Ojibwe *(this encompasses the land that Itasca is ‘on’)* in the Treaty of Washington, commonly known as the <u>1855 Treaty</u>. This treaty affirms the reserved rights doctrine and the inalienable rights of Ojibwe people to uphold their interminable relationship to the land. With affiliation to this and other academic institutions, it is our responsibility to acknowledge Native rights and the institutions’ history with them. We are committed to continue building relationships with the Ojibwe People through recognition, support, and to advocate for all Native American Nations. We strive to be good stewards of our place and privilege. *This land acknowledgement is a living document open to changes*.

## Acknowledgements

We thank SAK, Steve Knutie, Doug Thompson, Leroy, Paul for building nest boxes, and Itasca Biological Station (Emilie and Jonathan Schilling, Laura Domine, and Lesley Knoll) and UConn (OVPR office and Kent Holsinger) for logistical support during the 2020 pandemic. We also thank the following people for allowing us access to nest boxes on their property and their help in the field (in alphabetical order): Cheryl, Wade and Kelly Foy (Rock Creek General Store), Leroy, Lesley Knoll, Lake Itasca Pioneer Farmers, Tim Halberg, Brad and Kimberly Harsh, Helen and Ken Perry, Tim Qually, Greg and Vicki Qually, Sandy and Roger Smith, Doug and Dawn Thompson. We thank Bo Reese, Kendra Mass, and Lisa Stiepock at the University of Connecticut CORE facility for supplies and advice on molecular work and analyses.

Additionally, we thank Andrea Roth and Timothy Morre for providing statistical analysis advice and Daniel Bolnick and Jennifer Koop for comments on the manuscript. The work was funded by a Savaloja Research Grant from the Minnesota Ornithologists’ Union, the Research Grant from the North American Bluebird Society, EEB Research Grant from the UConn, Itasca-White Earth Research Fellowship Program, and UConn EEB summer funds to JG and start-up funds from the UConn to SAK. All applicable institutional guidelines for the care and use of animals were followed (UConn IACUC #18-005). Work was conducted under a SAK’s Federal Fisheries and Wildlife Scientific Collecting Permit (#MB11631D-0), Federal Master’s Banding Permit (#23623), and Minnesota Department of Natural Resources State Permit (#14878) and State Park Permit (#201936).

## Authors’ contribution statement

JG and SAK designed the study, JG, ST, and SAK collected the field data, JG and CG conducted the lab work, JG, CCT, and SAK conducted the statistical analyses, and JG, CCT, and SAK wrote the manuscript. All authors approved the final draft.

## References

Altermatt, F. & Ebert, D. 2008. Genetic diversity of Daphnia magna populations enhances resistance to parasites. Ecology Letters 11: 918–928.

Anderson, R.M. & May, R.M. 1982. Coevolution of Hosts and Parasites. Parasitology 85: 411–426.

Arct, A., Drobniak, S.M., Podmokła, E., Gustafson, L. & Cichoń, M. 2013. Benefits of extra-pair mating may depend on environmental conditions-an experimental study in the blue tit (Cyanistes caeruleus). Behavioral Ecology and Sociobiology 67: 1809–1815.

Blanchet, S., Rey, O., Berthier, P., Lek, S. & Loot, G. 2009. Evidence of parasite-mediated disruptive selection on genetic diversity in a wild fish population. Molecular Ecology 18: 1112–1123.

Boulinier, T., Sorci, G., Monnat, J.Y. & Danchin, E. 1997. Parent-offspring regression suggests heritable susceptibility to ectoparasites in a natural population of kittiwake Rissa tridactyla. Journal of Evolutionary Biology 10: 77–85.

Brinkhof, M.W.G., Heeb, P., Kölliker, M. & Richner, H. 1999. Immunocompetence of nestling great tits in relation to rearing environment and parentage. Proceedings of the Royal Society B: Biological Sciences 266: 2315–2322.

Chen, Y.H., Cheng, W.C., Yu, H.T. & Kam, Y.C. 2011. Genetic relationship between offspring and guardian adults of a rhacophorid frog and its care effort in response to paternal share. Behavioral Ecology and Sociobiology 65: 2329–2339.

Coster, G., Neve, L., Verhulst, S. & Lens, L. 2012. Maternal effects reduce oxidative stress in female nestlings under high parasite load. Journal of Avian Biology 43: 177–185.

Covarrubias-Pazaran G, Diaz-Garcia L, Schlautman B, Salazar W, Zalapa J. (2015) Fragman: An R package for fragment analysis. BMC Genetics 17(62):1–8 URL http://bmcgenet.biomedcentral.com/articles/10.1186/s12863-016-0365-6

DeSimone, J. G., Clotfelter, E. D., Black, E. C., & Knutie, S. A. 2018. Avoidance, tolerance, and resistance to ectoparasites in nestling and adult tree swallows. Journal of Avian Biology, 49: 1–12.

Ekroth, A.K.E., Rafaluk-Mohr, C. & King, K.C. 2019. Host genetic diversity limits parasite success beyond agricultural systems: a meta-analysis. Royal Society Publishing. B 286: 1–9.

Faircloth, B.C., Keller, G.P., Nairn, C.J., Palmer, W.E., Carroll, J.P. and Gowaty, P.A. 2006. Tetranucleotide microsatellite loci from eastern bluebirds Sialia sialis. Molecular Ecology Notes, 6: 646–649.

Ferree, E.D., Dickinson, J.L., Kleiber, D., Stern, C.A., Haydock, J., Stanback, M.T., Schmidt, V., Eisenberg, L. and Stolzenburg, C. 2008. Development and cross-species testing of western bluebird (Sialia mexicana) microsatellite primers. Molecular Ecology Resources, 8: 1348–1350.

Fox, J., & Weisberg, S. 2011. An R companion to applied regression (2nd ed.). Thousand Oaks, CA: Sage.

Giordano, M., Groothuis, T.G.G. & Tschirren, B. 2014. Interactions between prenatal maternal effects and posthatching conditions in a wild bird population. Behavioral Ecology 25: 1459–1466.

Grab, K.M., Hiller, B.J., Hurlbert, J.H., Ingram, M.E., Parker, A.B., Pokutnaya, D.Y., et al. 2019. Host tolerance and resistance to parasitic nest flies differs between two wild bird species. Ecology and Evolution 00: 1–12.

Griffith S.C., Stewart I.R.K., Dawson D.A., Owens I.P.F., Burke T. 1999. Contrasting levels of extra-pair paternity in mainland and island populations of the house sparrow (Passer domesticus): is there an “island effect”? Biological Journal of the Linnean Society 68:303–316

Hain, T.J.A. & Neff, B.D. 2007. Multiple paternity and kin recognition mechanisms in a guppy population. Molecular Ecology 16: 3938–3946.

Hasselquist, D. 2001. Social mating systems and extrapair fertilizations in passerine birds. Behavioral Ecology 12: 457–466.

Holt, R.D., Dobson, A.P., Begon, M., Bowers, R.G. & Schauber, E.M. 2003. Parasite establishment in host communities. Ecology Letters 6: 837–842.

Jennions, M.D. & Petrie, M. 2000. Why do females mate multiply? A review of the genetic benefits. Biological Reviews of the Cambridge Philosophical Society 75: 21–64.

King, M.O., Owen, J.P. & Schwabl, H.G. 2010. Are maternal antibodies really that important? patterns in the immunologic development of altricial passerine house sparrows (Passer domesticus). PLoS ONE 5: e9639.

Klein, S.L. 2000. The effects of hormones on sex differences in infection: From genes to behavior. Neuroscience and Biobehavioral Reviews 24: 627–638.

Knutie, S.A. 2020. Food supplementation affects gut microbiota and immunological resistance to parasites in a wild bird species. Journal of Applied Ecology. 00: 1–12.

Lively, C.M., de Roode, J.C., Duffy, M.A., Graham, A.L. & Koskella, B. 2014. Interesting open questions in disease ecology and evolution. American Naturalist 184: S1–S8.

Mateo & Holmes. 2004. Cross-fostering as a means to study kin recognition. Animal Behaviour 68: 1451–1459.

Murphy, W.G. 2014. The sex difference in haemoglobin levels in adults — Mechanisms, causes, and consequences. Blood Reviews, Elsevier B.V YBLRE-0032: 1–7

O’Brien, E.L. & Dawson, R.D. 2013. Nestling sex predicts susceptibility to parasitism and influences parasite population size within avian broods. Journal of Avian Biology 44: 226–234.

O’Brien, E.L. & Dawson, R.D. 2009. Palatability of passerines to parasites: Within-brood variation in nestling responses to experimental parasite removal and carotenoid supplementation. Oikos 118: 1743–1751.

Owen, J.P., Nelson, A.C. & Clayton, D.H. 2010. Ecological immunology of bird-ectoparasite systems. Trends in Parasitology 26: 530–539. Elsevier Ltd.

Palacios, M.G., Cunnick, J.E., Vleck, D. & Vleck, C.M. 2009. Ontogeny of innate and adaptive immune defense components in free-living tree swallows, Tachycineta bicolor. Developmental and Comparative Immunology 33: 456–463.

Penn, D.J., Damjanovich, K., Potts, W.K., Penn, D.J., Damjanovich, K. & Potts, W.K. 2002. MHC heterozygosity confers a selective advantage against multiple-strain infections. National Academy of Sciences 99: 11260–11264.

Pihlaja, M., Siitari, H. & Alatalo, R. v. 2006. Maternal antibodies in a wild altricial bird: Effects on offspring immunity, growth and survival. Journal of Animal Ecology 75: 1154–1164.

Råberg, L., Graham, A.L. & Read, A.F. 2009. Decomposing health: Tolerance and resistance to parasites in animals. Philosophical Transactions of the Royal Society B: Biological Sciences 364: 37–49.

Rolland, C., Macdonald, D.W., de Fraipont, M. & Berdoy, M. 2003. Free female choice in house mice: Leaving best for last. Behavior 140: 1371–1388.

Sabrosky, C.W., Bennett, G.F. & Whitworth, T.L. 1989. Bird blow flies (Protocalliphora) in North America (Diptera: Calliphoridae), with notes on the Palearctic species. Smithsonian Institution Press: 20–39.

Sommer, S. 2005. The importance of immune gene variability (MHC) in evolutionary ecology and conservation. Frontiers in Zoology 2: 1–18.

Stewart, S.L.M., Westneat, D.F. & Ritchison, G. 2009. Extra-pair paternity in eastern bluebirds: Effects of manipulated density and natural patterns of breeding synchrony. Behavioral Ecology and Sociobiology 64: 463–473.

Venables, W. N., & Ripley, B. D. 2002. Modern applied statistics with S (4th ed.). New York, NY: Springer: 183–300.

Winney, I., Nakagawa, S., Hsu, Y.H., Burke, T. & Schroeder, J. 2015. Troubleshooting the potential pitfalls of cross-fostering. Methods in Ecology and Evolution 6: 584–592.

Zuk, M. & McKean, K.A. 1996. Sex differences in parasite infections: Patterns and processes. International Journal for Parasitology 26: 1009–1024.

